# Redefining the role of Hypoxia-inducible factors (HIFs) in oxygen homeostasis

**DOI:** 10.1101/2024.03.15.585157

**Authors:** Clemente F. Arias, Francisco J. Acosta, Federica Bertocchini, Cristina Fernańdez-Arias

**Affiliations:** Grupo Interdisciplinar de Sistemas Complejos de Madrid (GISC), 28040 Madrid, Spain; Departamento de Ecología, Universidad Complutense de Madrid, 28040 Madrid, Spain; Plasticentropy, 51100 Reims, France; Departamento de Inmunología, Facultad de Medicina, Universidad Complutense de Madrid, 28040 Madrid, Spain

**Keywords:** Hypoxia-inducible factors (HIFs), metabolism, intracellular oxygen homeostasis, intracellular energy homeostasis, glycolysis, TCA cycle, fermentation, NAD+, redox balance, aerobic glycolysis, Warburg effect, pseudohypoxia

## Abstract

Hypoxia-inducible factors (HIFs) are key regulators of intracellular oxygen homeostasis. The marked increase in HIFs activity in hypoxia as compared to normoxia, together with their transcriptional control of primary metabolic pathways, motivated the widespread view of HIFs as responsible for the cell’s metabolic adaptation to hypoxic stress. In this work, we suggest that this prevailing model of HIFs regulation is misleading. We propose an alternative model focused on understanding the dynamics of HIFs’ activity within its physiological context. Our model suggests that HIFs would not respond to but rather prevent the onset of hypoxic stress by regulating the traffic of electrons between catabolic substrates and oxygen. The explanatory power of our approach is patent in its interpretation of the Warburg effect, the tendency of tumor cells to favor anaerobic metabolism over respiration, even in fully aerobic conditions. This puzzling behavior is currently considered as an anomalous metabolic deviation. Our model predicts the Warburg effect as the expected homeostatic response of tumor cells to the abnormal increase in metabolic demand that characterizes malignant phenotypes. This alternative perspective prompts a redefinition of HIFs’ function and underscores the need to explicitly consider the cell’s metabolic activity in understanding its responses to changes in oxygen availability.

## Introduction

Oxygen plays opposite physiological roles, being at once crucial for cellular respiration and a source of harmful free radicals that may compromise the cell’s function. This duality forces cells to maintain an exquisite oxygen balance, a task that relies to a great extent on Hypoxia-Inducible Factors (HIFs), transcription factors that control the expression of hundreds of genes involved in oxygen homeostasis both at cellular and systemic levels [1]. HIFs are heterodimers formed by two different subunits labeled alpha and beta. The former is marked as a target for ubiquitination and degradation through the proteasome pathway by HIF prolyl hydroxylase domain (PHDs) enzymes [2]. PHDs regulate the stability of HIF-*α* according to intracellular oxygen availability [2–6] by catalyzing the following reaction:

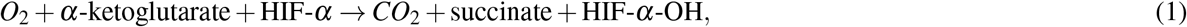

As a result of PHDs regulation, HIF-*α* expression is very high in cells cultured in 1% oxygen (a scenario typically referred to as hypoxia) and barely detectable under 20% oxygen (often labeled as normoxia) [7–9]. This pattern of expression is widely interpreted as giving rise to an intracellular sensor of hypoxia. HIFs would remain inactive in normoxia (Fig. 1.A), and they would only operate when extracellular oxygen levels are low (Fig. 1.B), triggering the expression of multiple target genes that adapt the cell’s metabolism to oxygen deprivation [10].

**Figure 1.**
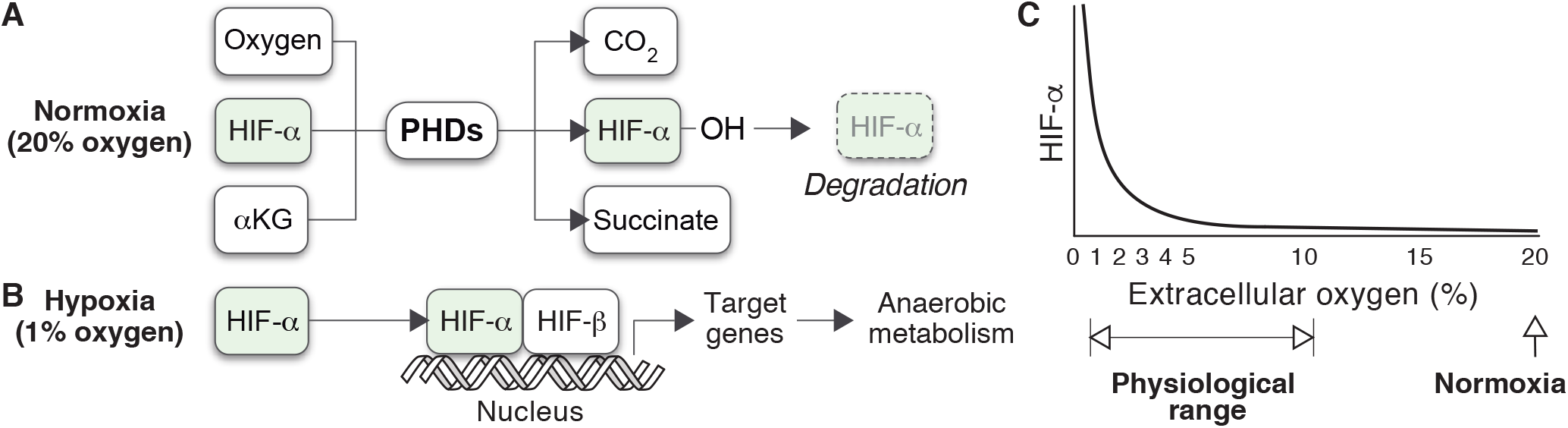
Prevailing model of HIFs regulation. A,B) HIF-*α* is currently assumed to follow a bimodal mode of expression, determined by extracellular oxygen tensions. In normoxia, the rate of PHD-mediated degradation suffices to maintain HIF-*α* expression at residual levels (A). The absence of oxygen restricts the activity of the PHDs, leading to the stabilization of HIF-*α* and enabling the activation of target genes (B). C) Dependence of HIF-*α* expression on extracellular oxygen tensions as observed in HeLa cells.

Normoxia and hypoxia have become standard experimental conditions in the field since the seminal studies that led to the discovery of HIFs [7]. Now, oxygen tensions within the human body broadly vary between less than 1% and 11% [11–13], with most cell types encountering oxygen tensions below 6%. This is the case in organs such as the liver or the uterus (4-6.5% and 2-2.5% oxygen respectively) [12], and even the brain (4-6.4% oxygen) [14]. Remarkably, partial pressures of oxygen in the large intestine can be as low as 0.4-1.5% [12].

Therefore, normoxia is well above the physiological range normally found by cells inside the organism. In fact, 20% oxygen is close to the most hyperoxic environment that cells can experience in natural circumstances on Earth. Nevertheless, normoxia has traditionally been used as the reference status for oxygen homeostasis. Accordingly, the absence of HIF-*α* is often interpreted as the normal, default state of the cell, and its expression is typically described as an active cellular response to low oxygen availability [15–19].

Empirical evidence challenges this view of HIFs activity. An early study involving HeLa cells cultured under various extracellular oxygen conditions found that HIF-*α* expression was notably high within the oxygen range typical of body tissues [20, 21] (Fig. 1.C). HIF-*α* levels only became negligible when the partial pressure of oxygen exceeded the physiological threshold typically encountered by cells (Fig. 1.C).

The expression of HIF-*α* in a wide range of oxygen tensions underlines a critical limitation of the prevailing model of HIFs regulation. This model accounts for the difference in HIF-*α* levels under normoxia and hypoxia, which are two extreme situations of extracellular oxygen availability. However, it does explain the expression of HIF-*α* at intermediate oxygen levels, which correspond to physiological conditions. If we accept that HIF-*α* is a sign of oxygen deficit [22, 23], cells from tissues below 10% oxygen would undergo persistent hypoxic stress in the organism (Fig. 1.C).

In this work, we suggest that the dichotomy between normoxia and hypoxia is artificial and hinders our understanding of intracellular oxygen homeostasis. HIFs regulation should be interpreted within its physiological context, which implies that intermediate oxygen levels should be explicitly considered. To explain the homeostatic function of HIFs within the cell, we formulate a conceptual model that integrates key metabolic processes regulated by HIFs, such as fermentation or glycolysis. Then, we use this model to analyze the impact of these processes on intracellular oxygen homeostasis.

Our model suggests that the homeostatic role of HIFs should be redefined. HIFs would regulate the transit of electrons from catabolic substrates to oxygen, preventing the saturation of the cellular pathways of energy generation. From this approach, HIFs would not coordinate the adaptation of the cell’s metabolism to hypoxic stress. Instead, they would prevent the onset of hypoxic stress in the first place.

According to our model, the pattern of HIF-*α* expression observed in HeLa cells (Fig. 1.C) is the expected outcome of Reaction 1 when it is considered within its metabolic context. As a result, HIFs would not exhibit a bimodal pattern of activity. Varying HIF-*α* levels would be necessary to adjust the fraction of high-energy electrons generated in metabolic pathways that are eventually transferred to oxygen depending on cellular oxygen uptake.

Contrary to the prevailing paradigm, our model also suggests that homeostatic imbalance cannot only arise from a drop in extracellular oxygen availability. An increase in the supply of catabolic substrates, which are the source of high-energy electrons eventually used to generate energy, may also cause a cellular deficiency of oxygen. This approach stresses the role of major pathways of energy generation in the cellular demand for oxygen, a key aspect of oxygen homeostasis that is not explicitly present in the prevailing model of HIFs regulation (Figs. 1.A,B).

The lack of an appropriate metabolic context in the current explanation of HIFs regulation has led to profound misconceptions about the cell’s use of oxygen. A notable example is the predominance of anaerobic metabolism observed in tumor cells under fully aerobic conditions, a phenomenon known as the Warburg effect (WE) [24, 25]. Within the current paradigm, the prevalence of fermentation over aerobic respiration is perceived as an abnormal metabolic response to the constraints imposed by the tumor environment [26]. The WE would be a hallmark of tumor cells and, consequently, a deviation from normal cellular homeostasis [26]. Our model suggests that the WE is the expected consequence of HIF-*α* upregulation in tumor cells, which in turn would be the normal, physiological response to the anomalous increase in metabolic demand that defines malignant phenotypes. From this viewpoint, the ability to maintain oxygen homeostasis in tumors would be an outstanding manifestation of the functional flexibility conferred by HIFs regulation.

This work contributes to a better understanding of HIFs’ pivotal role in cellular oxygen homeostasis. Our results broaden our comprehension of how HIFs operate in their intracellular physiological context and underscore the role of cell’s metabolism in oxygen homeostasis. The insights provided by our model point to common underlying mechanisms operating in a wide array of conditions, ranging from hypoxia to disorders such as tumors or diabetes. The new understanding arising from this approach has profound implications for future studies on cellular functions and may open new avenues for innovative therapeutic interventions.

## Results

### The physiological context of HIFs regulation

The cycle of NAD+ (nicotinamide adenine dinucleotide) and NADH (the reduced form of NAD+) is central to primary metabolic processes involved in cellular energy production, including glycolysis, fermentation, and respiration. It plays a crucial role in the transfer of high-energy electrons obtained from the catabolic breakdown of glucose and other substrates to the electron transport chain (ETC) and, subsequently, to oxygen (Fig. 2).

**Figure 2.**
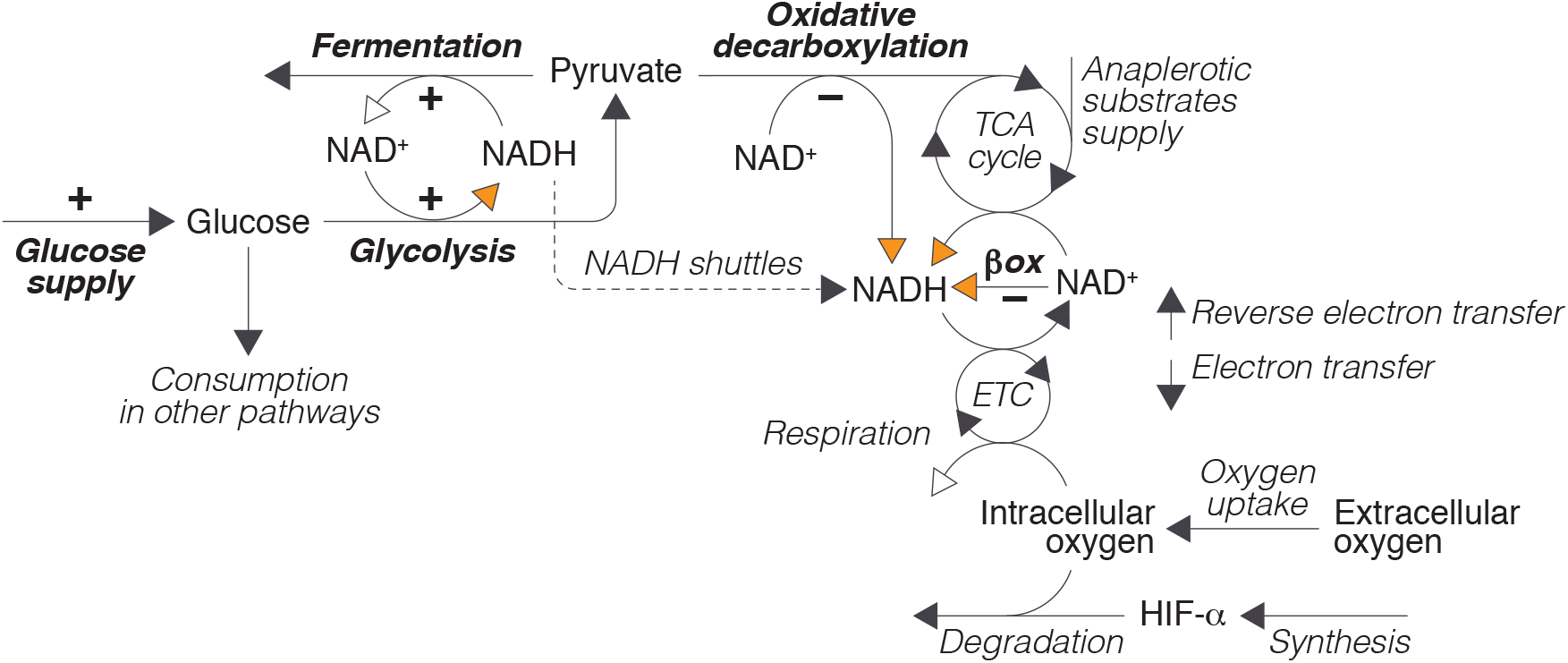
Conceptual model of the physiological context of HIFs activity. Glucose (obtained from extracellular uptake or from intracellular reserves) can be used in different cellular processes. In glycolysis, it is partially broken down into pyruvate to produce ATP. Pyruvate can then be converted into lactate by the cytoplasmic enzyme lactate dehydrogenase A (LDHA), or, alternatively, enter the tricarboxylic acid (TCA) cycle after being converted into acetyl-CoA through oxidative decarboxylation by the pyruvate dehydrogenase (PDH) enzymatic complex. High-energy electrons produced during glycolysis or from the breakdown of carbon compounds (pyruvate and other anaplerotic substrates) during the TCA cycle and the *β* − oxidation of fatty acids are eventually transferred to the ETC and, from there, to oxygen. This process is an integral part of oxidative phosphorylation, which drives the production of ATP in aerobic respiration, and constitutes the primary cause of oxygen consumption in the cell. Oxygen also participates in HIF-*α* degradation by PHDs according to Reaction 1. Orange and white arrows represent the entry and exit of electrons into and out of the NAD+/NADH cycle respectively. The processes regulated by HIFs are indicated in bold and the signs indicate if HIFs regulation is positive or negative.

The transit of electrons through the NAD+/NADH cycle is intimately linked to the cellular utilization of pyruvate, a key metabolite located at the intersection of the main pathways of energy generation in the cell. Pyruvate serves as a substrate for both the TCA cycle and lactic fermentation, so it defines a pivotal branching point where the cell’s metabolism can diverge into either aerobic or anaerobic pathways (see Fig. 2).

The oxidative decarboxylation of pyruvate and its entry into the TCA cycle represent further steps in the oxidation of glucose initiated in glycolysis, and supply additional high-energy electrons to the NAD+/NADH cycle that are eventually transferred to oxygen (Fig. 2). In contrast, fermentation hinders the full breakdown of glucose in the TCA cycle by diverting pyruvate towards lactate production, thus bypassing its entry into the mitochondria. In addition, electrons generated in glycolysis are used during fermentation to reduce pyruvate instead of being shuttled into the mitochondria to generate energy in aerobic respiration, which further decreases the production of ATP from the catabolism of glucose. Therefore, the fraction of the electrons obtained from the catabolism of glucose that are eventually transferred to oxygen largely depends on whether pyruvate is used in fermentation or in the TCA cycle (Fig. 2).

This metabolic choice is a primary target of HIFs regulation (Fig. 2). HIFs control pyruvate production both directly, by upregulating glycolytic enzymes [18, 27–29], and indirectly, by stimulating the synthesis of glucose transporters (GLUT1 and GLUT3) that facilitate the transport of glucose across the cell membrane, increasing the availability of glucose for glycolysis [27, 27, 30–32].

HIFs also regulate pyruvate consumption, promoting its use in lactic fermentation at the expense of aerobic respiration. On one hand, they accelerate pyruvate conversion to lactate by upregulating LDHA expression [19, 33]. Additionally, they inhibit respiration by upregulating PDH kinase 1 (PDK1) [34, 35], an enzyme that phosphorylates and inactivates PDH, reducing the conversion of pyruvate to acetyl-CoA [36] and its use in the TCA cycle.

Besides regulating the intracellular metabolism of pyruvate, HIFs also inhibit the *β* -oxidation of fatty acids, which represents another major input of electrons into the system, as well as a source of acetyl-CoA that can be used as substrate to feed the TCA cycle [37].

In summary, the regulatory activity of HIFs is embedded in a complex intracellular metabolic network that defines the cellular use of pyruvate and the transit of electrons through the NAD+/NADH cycle. We suggest the conceptual model shown in Fig. 2 provides a functional description of this network. We will use a mathematical formulation of this conceptual model (see Supplementary Material) to gain insight into the dynamics of HIFs regulation within its physiological context. To that end, we will compare the behavior of the system with and without HIFs activity. The rationale for this approach is that, in the absence of HIFs control, the system should easily undergo homeostatic dysregulation. HIFs are expected to change the default system’s dynamics to prevent the onset of homeostatic imbalance. By analyzing these HIFs-induced changes it is possible to discern the consequences of HIFs regulation, which is key to a better understanding of oxygen homeostasis.

### Redefining the homeostatic role of HIFs

HIFs are widely defined as intracellular sensors of hypoxia. However, the term “hypoxia” is ambiguous as it indistinctly used in the literature to designate low oxygen supply and oxygen deficit. Both concepts are not equivalent though, since the former refers to absolute extracellular oxygen levels (typically 1% atmospheric oxygen), whereas the latter is a relative condition that denotes an imbalance between oxygen needs and availability. Taking both concepts as synonyms implicitly assumes that low oxygen levels necessarily entail a cellular deficit and, conversely, that this deficit is impossible in normoxia (Figs. 1.A,B).

This assumption, which lays at the basis of the current paradigm of oxygen homeostasis, is questionable. To clarify this point, let us analyze the dynamics of the ETC and the NAD+/NADH cycle in the absence of homeostatic regulation (see Supplementary Material for details). Without HIFs control, the ETC is saturated with electrons when oxygen uptake is too low relative to glucose supply (Fig 3.A). In this scenario, electrons move from the ETC back to NAD+ through reverse electron transfer [38], saturating the NAD+/NADH cycle (Fig 3.A). This saturation propagates backward across the system, causing the concomitant accumulation of intermediate metabolites such as acetyl-CoA (Fig. 3.B).

**Figure 3.**
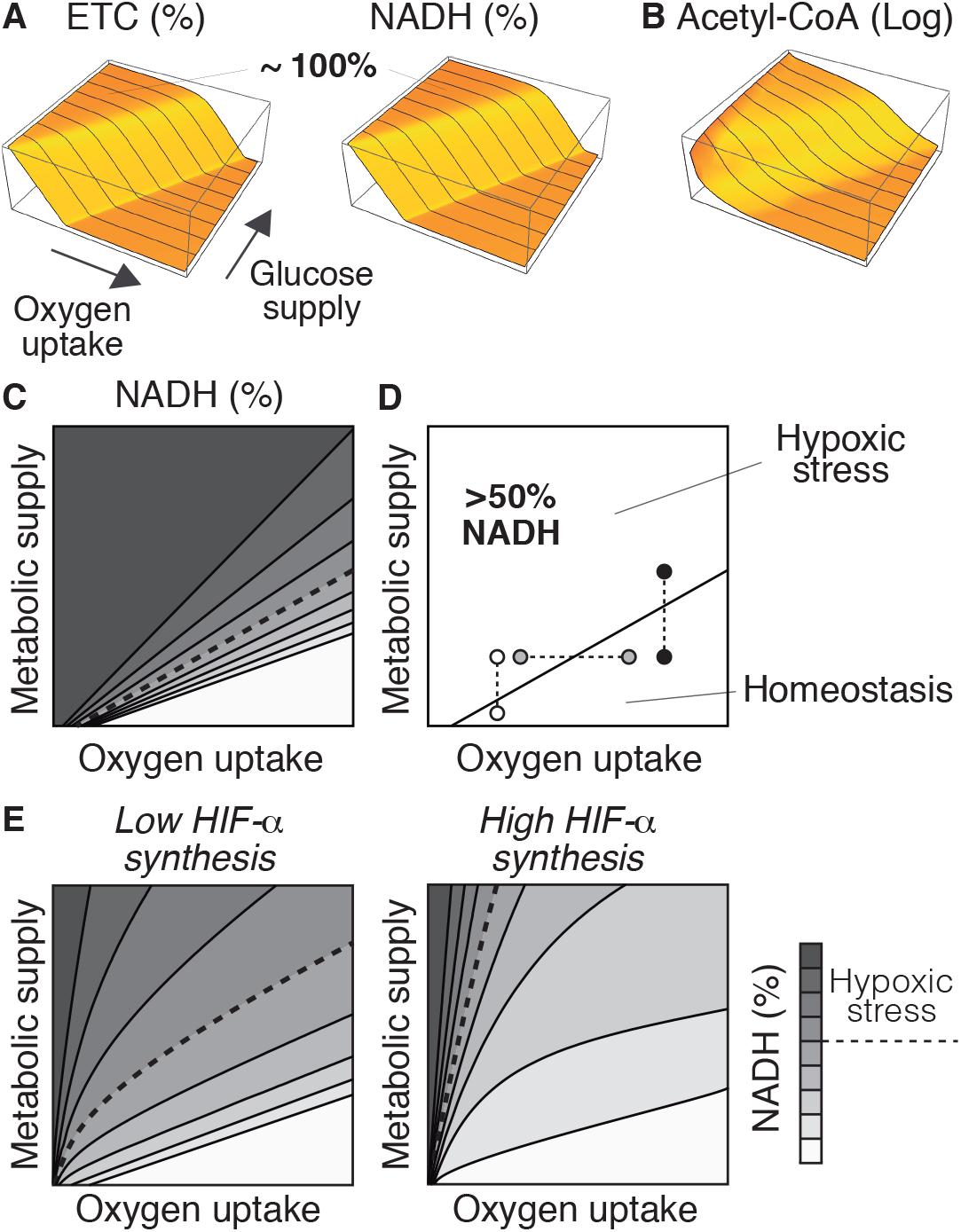
Definition of hypoxia and hyperoxia in relation to the saturation of the NAD+/NADH cycle. A) Saturation of the ETC and the NAD+/NADH cycle as a function of oxygen uptake and glucose supply. B) Accumulation of acetyl-CoA in the system as a function of oxygen uptake and glucose supply. C) Contour plot of the saturation of the NAD+/NADH cycle (defined as %NADH) as a function of oxygen uptake and metabolic supply (defined as the input of glucose, fatty acids, and anaplerotic substrates for the TCA cycle). D) Values above 50% NADH correspond to homeostatic stress. This threshold delimits the homeostatic range of the system, i.e. the values of oxygen and metabolic input for which the system is in homeostatic balance. E,F) Variations in the rate of HIF-*α* synthesis greatly affect the homeostatic range of the system. Combination of inputs that homeostatic balance may result in hypoxic stress when IF-*α* synthesis decreases. The choice of 50% NADH is arbitrarily chosen and the same arguments apply for any other threshold value.

We can infer from these results that the accumulation of NADH is an indicator of oxygen deficit. Within the cell, an excess of NADH exacerbates the production of reactive oxygen species, compromising the cell’s functionality [39–41] and contributing to pathological conditions at both the cellular and systemic levels [42–44]. For this reason, we will refer to the accumulation of NADH as hypoxic stress (Fig. 3.C), and exclusively use the term ‘hypoxia’ to denote low oxygen levels.

From our approach, hypoxic stress does not only depend on oxygen availability but also on the metabolic activity of the cell. Thus, hypoxia does not necessarily entail hypoxic stress (white circles in Fig. 3.D) and, reciprocally, high oxygen levels do not necessarily prevent it (black circles in Fig. 3.D). For a given value of oxygen uptake, an increase in metabolic supply may cause a deficit of oxygen (white and black squares in Fig. 3.D). Analogously, for each particular value of metabolic supply, the system can undergo hypoxic stress if oxygen uptake decreases (gray circles in Fig. 3.D).

In agreement with these results, intracellular NAD+ exceeds NADH by one or two orders of magnitude under normal circumstances [45] but the balance may shift toward NADH in hypoxia [46–49] or when glucose levels are abnormally high (as occurs during diabetes) [41, 50–53], a phenomenon that has been termed pseudohypoxia [50].

Taking hypoxia and hypoxic stress as synonyms has led to deep-rooted misunderstandings regarding the role of HIFs in intracellular oxygen homeostasis. A notable example of this is pseudohypoxia. According to the prevailing paradigm of oxygen homeostasis, the intracellular accumulation of NADH under hypoxia is easily explained as a consequence of oxygen deficit [46–49, 54]. However, in pseudohypoxia, the accumulation of NADH takes place in normal oxygen conditions, so it is viewed as unrelated to oxygen homeostasis [41, 50–53].

Furthermore, since oxygen deficit is considered incompatible with normal oxygen levels, the upregulation of HIF-*α* observed in pseudohypoxic cells is perceived as puzzling [51, 55, 56]. This has motivated the search for alternative, oxygen-independent mechanisms of HIF-*α* stabilization during pseudohypoxia [56, 57]. From this approach, the degradation of HIF-*α* by PHDs would depend on oxygen levels in hypoxia, but it would be governed by different mechanisms in pseudohypoxia.

We propose that hypoxia and pseudohypoxia are particular cases of hypoxic stress. Both conditions would be caused by an imbalance between oxygen uptake and metabolic supply, leading to the accumulation of NADH in the system. In hypoxia, the imbalance would arise from a drop in oxygen availability, whereas in pseudohypoxia it would result from an increase in metabolic supply. Despite their apparent differences, the expression of HIF-*α* would respond to the same underlying principle in both cases.

As for the consequences of HIF-*α* expression, our model suggests that it reduces the range of conditions that lead to hypoxic stress (Fig. 3.E). HIFs regulation prevents the accumulation of NADH under combinations of oxygen uptake and metabolic supply that would cause homeostatic imbalance without HIFs. This protective effect of HIFs increases with the rate of HIF-*α* synthesis (Fig. 3.E).

In this regard, it is important to stress that the prevailing model of HIFs regulation is focused on the degradation of HIF-*α* by PDHs, and does not explicitly consider HIF-*α* production (Figs. 1.A,B). However, the *HIF1A* gene is subjected to complex transcriptional regulation [10, 58], which suggests that cells might adapt the rate of HIF-*α* synthesis depending on the circumstances. This would open novel ways to modulate HIFs control of intracellular oxygen homeostasis (Fig. 3.E). We will return to this point below.

Based on the previous results, we suggest that HIFs regulate the transit of electrons through the NAD+/NADH cycle to prevent the saturation of the system (Figs. 3.C,E). In the following sections, we will show that this regulation would be necessary to maintain an adequate balance between NAD+ and NADH in a wide range of metabolic conditions.

### HIFs response to changes in oxygen availability

In this section, we will assume a fixed value of metabolic supply (glucose, fatty acids, and anaplerotic substrates) and analyze the HIFs response to changes in oxygen uptake. According to our model (see Supplementary Material), HIF-*α* progressively decreases as extracellular oxygen tensions increase, and only becomes negligible at high oxygen tensions (Fig. 4.A). This result predicts the pattern of HIF-*α* expression observed in HeLa cells at physiological oxygen levels [20] (see Fig. 1.C). The presence of HIF-*α* in a wide range of oxygen levels suggests that target genes are not transcribed in an all-or-nothing manner, as assumed by the current model of HIFs regulation. Instead, the degree of gene expression could vary depending on HIF-*α* levels, adjusting the reaction rates of HIFs regulated processes according to oxygen levels (Figs. 3.B-D).

**Figure 4.**
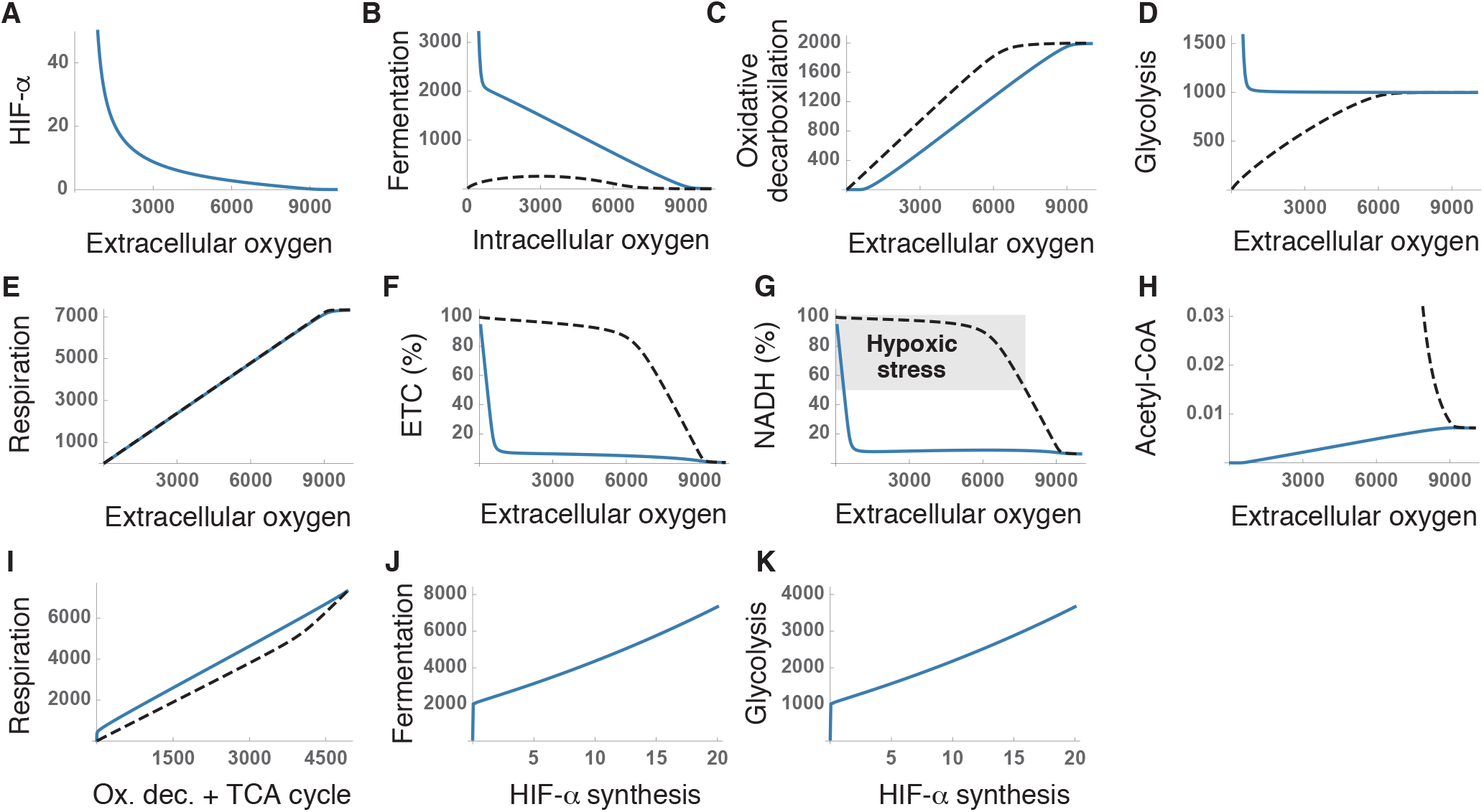
Homeostatic response to changes in extracellular oxygen tensions. A) Variation of HIF-*α* levels with extracellular oxygen tensions. B-D) Response of the metabolic pathways controlled by HIFs to changes in oxygen availability with and without regulation. E) Effect of extracellular oxygen tension on aerobic respiration. F,G) Saturation of the ETC (F) and the NAD+/NADH cycle (G) with electrons for different extracellular oxygen tensions. H) Accumulation of acetyl-CoA in the system. I) Aerobic respiration as a function of the summed rates of oxidative decarboxylation and the TCA cycle. J,K) Effect of the rate of HIF-*α* synthesis on fermentation (J) and glycolysis (K). These results correspond to numerical simulations of our model for a fixed input of glucose, fatty acids, and anaplerotic substrates (see Supplementary Material). Axis units are arbitrary and the values are shown for illustrative purposes. Solid and dashed lines correspond to simulations with and without HIFs regulation respectively.

The homeostatic role of HIFs is widely interpreted as switching the cell’s metabolism toward anaerobic pathways, sustaining ATP production when oxygen becomes a limiting factor [10, 28, 29, 35, 59–62]. In our model, aerobic respiration is constrained by oxygen availability and is greatly reduced as oxygen availability decreases (Fig. 4.E). The upregulation of glucose uptake and glycolysis (Fig. 4.D), together with the inhibition of oxidative decarboxylation (Fig. 4.C) certainly fosters anaerobic metabolism at low oxygen levels. However, this explanation does not easily account for the marked upregulation of fermentation, or the substantial expression of HIF-*α* observed at intermediate oxygen levels (Figs. 4.A,B).

These results can be better interpreted under the assumption that HIFs are intended to regulate the transit of electrons through the NAD+/NADH cycle. HIFs’ regulation prevents the saturation of the ETC (Fig. 4.F) and maintains adequate NADH levels across a broad range of oxygen tensions (Fig. 4.G), additionally preventing the accumulation of intermediate metabolites, such as acetyl-CoA (Fig. 4.H). In the absence of HIFs activity, the system easily becomes saturated (Figs. 4.F,G), highlighting the essential role of HIFs in the transit of electrons across the system. Furthermore, the upregulation of fermentation by HIFs enhances the efficiency of aerobic respiration. Sustaining a given respiration rate necessitates a greater input of resources (i.e. higher rates of oxidative decarboxylation and TCA cycle) when HIFs are not active (Fig. 4.I)

In our model, the rate of HIF-*α* synthesis emerges as a key regulatory mechanism of critical relevance for the cell’s metabolism. We have seen that, in low oxygen regimes, HIFs upregulate fermentation and glycolysis (Figs. 4.D,E). Raising the rate of HIF-*α* synthesis greatly magnifies this effect (Figs. 4.J,K), which suggests that modulating HIF-*α* production could allow cells to fine-tune the production of ATP and lactate under oxygen deprivation.

In summary, our model suggests that HIFs could regulate the transit of electrons through the NAD+/NADH cycle as a function of oxygen availability, preventing the onset of hypoxic stress in a wide range of intracellular oxygen tensions. In the following section, we will explore how HIFs could also adapt the cell’s function to changes in metabolic supply. We will show that the efficiency of this homeostatic mechanism would be particularly evident for cells that exhibit an intense metabolic activity, as is often the case of tumor cells.

### The Warburg effect: normal oxygen homeostasis in tumor cells

The Warburg effect (WE) is the tendency of cancer cells to generate energy through the combination of glycolysis and fermentation (known as aerobic glycolysis or AG) instead of using the TCA cycle and oxidative phosphorylation, even in normoxia [56, 63–65]. The WE constitutes a metabolic signature of 70–80% of human cancers [66], with the fraction of ATP produced through glycolysis varying from less than 1% to over 60% [65, 67]. Ever since its discovery, the WE has been considered a puzzling phenomenon. Although some tumors show mitochondrial dysfunction, this is not the case in most cancer cells, which can normally produce ATP through oxidative phosphorylation [68–71]. Therefore, since aerobic respiration is more efficient in terms of energy production, tumor cells seem to incur an unnecessary metabolic cost by prioritizing AG when oxygen is not a limiting factor [60, 68, 72].

To account for this paradoxical behavior, it has been suggested that the WE would confer some type of competitive advantage to cancer cells over normal tissues, thus compensating for its apparent metabolic cost [26, 60, 73, 74]. Abnormal metabolic reprogramming would be key for tumor cells to obtain the energy they need to sustain intensive cell proliferation [26, 56, 66, 69, 73], constituting a hallmark of invasive cancers and one of the first steps in the development of malignant phenotypes [26, 66, 75].

Our model challenges this view of the WE as a hallmark of cancer and suggests that it is the expected physiological response to the marked increase in the uptake of nutrients, particularly glucose, that characterizes tumor cells [25, 69, 76]. According to our model (see Supplementary Material), raising metabolic supply (defined as the input of catabolic substrates into the system) should upregulate HIF-*α* expression in tumor cells, even under normal oxygen conditions (Fig. 5.A). It is important to stress that, in this case, HIF-*α* upregulation would not respond to changes in oxygen levels but exclusively to changes in the supply of catabolic substrates. Raising HIFs activity would maintain a fluid transit of electrons through the NAD+/NADH cycle when metabolic demand increases, preventing the onset of hypoxic stress (Fig. 5.B).

**Figure 5.**
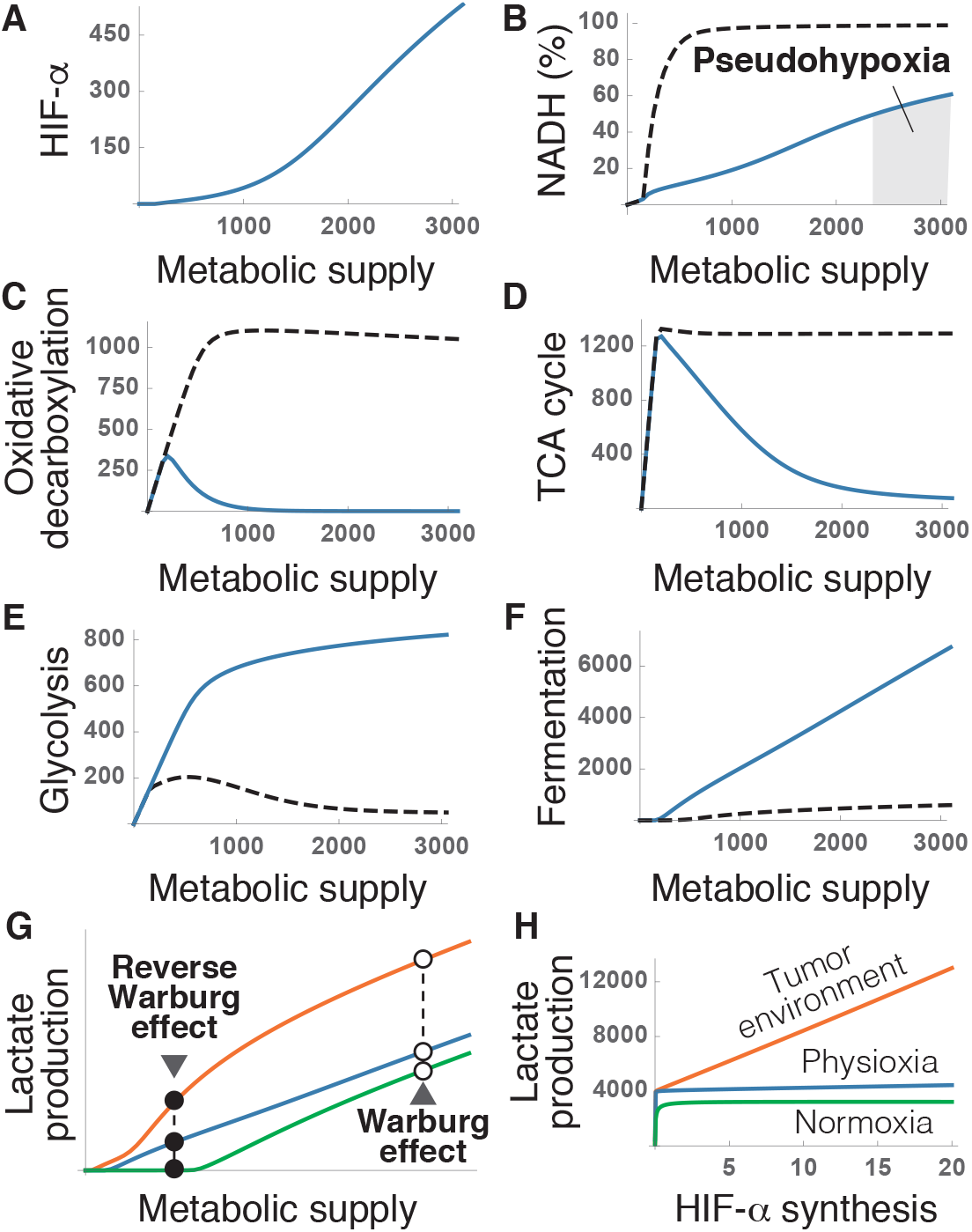
Homeostatic response to changes in metabolic supply. A) Dependence of HIF-*α* expression on the input of glucose, fatty acids, and anaplerotic substrates. B) Saturation of the NAD+/NADH cycle with electrons as a function of metabolic supply. The shaded region indicates saturations above 50% threshold that defines hypoxic stress. C,D) Downregulation of oxidative decarboxylation (C) and the TCA cycle (D) by HIFs metabolic supply. E,F) Upregulation of glycolysis (E) and fermentation (F) by HIFs. G) Lactate production as a function of metabolic supply for three different conditions of oxygen availability. These results correspond to numerical simulations of our model for a fixed value of oxygen uptake (see Supplementary Material). Axis units are arbitrary and the values are shown for illustrative purposes. Solid and dashed lines correspond to simulations with and without HIFs regulation respectively.

To that end, HIFs would induce a sharp reduction in the rates of oxidative decarboxylation and the TCA cycle (Figs. 5.C,D), and a concomitant boost of glycolysis and fermentation (Figs. 5.E,F). Such metabolic adaptations, occuring at normal oxygen levels, are the defining feature of the WE. According to our model, they would not exist without HIFs regulation (Figs. 5.B-F). The excess of glucose supply would also lead to the accumulation of NADH observed in normal cells during pseudohypoxia (Fig. 5.B).

In agreement with the previous results, tumors exhibit high levels of HIF-*α* expression, even in normoxia [26, 56, 66, 77], which leads to the upregulation of HIFs target genes when oxygen is amply available [64, 78]. These observations do not fit easily in the current paradigm of oxygen homeostasis (see Figs. 1.A,B), so they have been typically interpreted as anomalous features of tumor phenotypes [5, 66, 79, 80].

Our results suggest that increased HIF-*α* expression in tumors is not abnormal. The ability to proliferate without control is a distinctive trait of cancer cells that entails a significant increase in the supply of metabolic resources [25, 69, 76, 81]. According to our model, this behavior would increase the risk of hypoxic stress Fig. 5.B).The upregulation of HIF-*α* would be a physiological adaptation to ensure the viability of tumor cells in these circumstances. This would explain why experimentally increasing HIF-*α* expression fosters tumor growth, whereas inhibiting of HIFs activity hinders tumor progression [13, 79].

From this approach, the WE would be caused by the same homeostatic mechanisms that operate in healthy cells under normal circumstances. In fact, normal cells continuously produce lactate even under fully aerobic conditions [13, 33, 61, 82–84]. Both glucose uptake and lactate production increase in normal cells when they are stimulated to proliferate, a behavior that closely resembles the WE described in tumors [25]. Within our model, the increase in the supply of metabolic substrates caused by a surge in energy demand would upregulate HIF-*α* and increase lactate production both in normal and tumor cells (Fig. 5.G). This reaction would be exacerbated when oxygen availability is reduced (Fig. 5.G).

HIFs upregulation in hypoxia would therefore explain the observed increase in lactate production by normal cells within tumor environments, a phenomenon that has been labeled as the Reverse WE [65, 85, 86]. This effect could be further amplified by augmenting HIF-*α* synthesis. Changing this variable would have no apparent consequences at physiological and hyperoxic conditions, but it would dramatically increase the homeostatic production of lactate in hypoxia (Fig. 5.H). Modulating the rate of HIF-*α* synthesis would therefore provide an additional regulatory mechanism to increase glycolysis and lactate production in low oxygen conditions.

## Discussion

In this work, we redefine the role of HIFs in intracellular oxygen homeostasis. To do that, we formulate a conceptual model of the physiological context in which HIFs operate inside the cell (Fig. 2). A mathematical version of this model suggests that, in the absence of HIFs, the NAD+/NADH cycle becomes easily saturated with electrons when oxygen levels are low or metabolic supply (defined as the intracellular availability of catabolic substrates) is high. HIFs regulation prevents such imbalance, allowing the system to operate in a wide range of conditions. Based on these results, we suggest that HIFs are intended to regulate the transit of electrons through the NAD+/NADH cycle, ensuring a fluid electron transfer from catabolic substrates to oxygen.

The diagram shown in Fig. 2 represents a minimal, self-contained model system linking pyruvate metabolism, the NAD+/NADH cycle, and oxygen homeostasis. The scope and applicability of this model are constrained by its considerable degree of simplification (see Supplementary Material). For instance, it does not consider other electron carriers, such as FADH2, or the existence of additional regulatory elements, such as sirtuins, that also control the NAD+/NADH cycle [87]. Moreover, it does not consider other regulatory effects of HIFs, such as the control of mitochondrial mass [10] or the role of metabolites such as *α*-ketoglutarate, succinate, or fumarate in the activity of PHDs [88]. More importantly, it isolates a metabolic network that lies at the core of the cellular function and interacts with a myriad of other intracellular processes.

Nevertheless, this simplified model provides a suitable functional context in which to explore the dynamics of HIFs regulation. Its simplicity is key to identifying the functional consequences of these dynamics, something that would be complicated with more intricate models. The mathematical translation of this conceptual model is intended as a toy model to simulate its dynamics, and not as an accurate quantitative description of the behavior of the corresponding intracellular processes. Still, it provides valuable insight that can be extrapolated to the function of HIFs within the cell, an insight that would be hardly attainable without resorting to mathematical modeling.

Our model makes a series of predictions about oxygen homeostasis that have been experimentally validated in the literature: i) cells should normally express HIF-*α* in a wide range of oxygen tensions and not only in hypoxia, as observed in HeLa cells [20]; ii) an increase in glucose supply should raise HIF-*α* expression and cause the accumulation of NADH (pseudohypoxia) [41, 50–53]; iii) cancer cells should upregulate HIF-*α* and aerobic glycolysis, even in normoxia (WE) [26, 56, 63–65, 78]; and iv) healthy cells should normally produce lactate in fully aerobic conditions, and much more so in hypoxia (Reverse WE) [13, 33, 61, 82–84].

All these observations can therefore be explained as consequences of the functional relationships shown in Fig. 2. In contrast, some of them (such as HIF-*α* expression in pseudohypoxia and in tumors under normal oxygen conditions) are contradictory with the prevailing model of HIFs regulation, and require supplementary *a*d hoc explanations [56, 79]. In our opinion, these contradictions reflect critical limitations of the current paradigm, which result from a series of postulates that have been taken for granted in the field and whose validity is questionable.

First, it has long been thought that, since aerobic respiration is more efficient than AG, cells should only resort to fermentation when oxygen is not available, either to sustain energy production [10, 28, 29, 35, 59–62] or to prevent excess generation of ROS in the mitochondria under hypoxic conditions [13, 84]. This assumption does not take into account that the capacity of the NAD+/NADH cycle to transport electrons constrains the efficiency of respiration, or that this efficiency critically depends on other intracellular processes. HIFs control of fermentation would allow the cell to regulate the entry of pyruvate into the TCA cycle, thereby adjusting the cellular demand for oxygen to its extracellular availability. According to our model, this mode of action would prevent the saturation of the NAD+/NADH cycle, maintaining the energy yield of respiration when oxygen availability decreases or the cell’s metabolic demand increases.

A second postulate widely assumed in the literature is that hypoxia is equivalent to hypoxic stress. This approach has biased the view of HIFs as exclusively responding to changes in oxygen levels, neglecting the role of cellular metabolism in the onset of hypoxic stress. Changes in energy demand and, consequently, in the supply of substrates for catabolic pathways, are as common as changes in oxygen levels. According to our model, both variables participate in defining the cell’s demand for oxygen, and both control HIFs activity.

Finally, it has long been assumed that normoxia and hypoxia are appropriate experimental conditions to understand oxygen homeostasis. However, they represent two extremes of a continuum of oxygen availability. Our model shows that HIFs regulation is key to maintain the flow of electrons through the NAD+/NADH cycle at intermediate oxygen tensions, which coincide with the conditions normally found by cells in their physiological context. The variation in HIF-*α* expression with oxygen tensions observed in HeLa cells would be the expected physiological manifestation of HIFs regulation.

The previous assumptions have critically conditioned the interpretation of empirical evidence, leading to profound misconceptions about HIFs function. In particular, the description of HIF-*α* as a sensor of hypoxia is a phenomenological model based on differences between cells in normoxia and hypoxia (see Fig. 1). This model has motivated the view of HIFs as orchestrating a cellular response to hypoxic stress [33, 88]. In our opinion, such a HIFs-mediated stress response would provide a deficient homeostatic adaptation.

It is important to recall that HIFs regulation involves the transcription of target genes, a process that may take hours. According to the current paradigm, in case of hypoxia, HIFs should first accumulate in the cell’s cytoplasm and then migrate to the nucleus, where they would activate the transcription of target genes, and the subsequent synthesis of target proteins. Only at this point would the cell’s metabolism be ready to operate in hypoxia. This mechanism of action would entail a marked delay between stress sensing and the cell’s response. Furthermore, this response would rely on demanding cellular activities, such as gene transcription and protein synthesis, operating under stress conditions where oxygen, and therefore energy production, are hindered. This suggests that additional, faster regulatory mechanisms must be responsible for adapting the cell’s function to sudden changes in conditions until an alternative, appropriate metabolic configuration is established by HIFs [89, 90].

HIFs transcriptional control can be better understood as defining the default, long-term organization of the cell’s metabolism according to oxygen availability. This would be achieved by setting the reaction rate of processes that play key roles in the cell’s use of oxygen, such as fermentation, glycolysis, or beta-oxidation (Fig. 2).

In this regard, the name of Hypoxia-inducible factors is an unfortunate choice. The focus on hypoxia has motivated that phenomena such as NADH accumulation or HIF-*α* expression during pseudohypoxia or the WE are perceived as paradoxical, since they occur under normal oxygen conditions. For this reason, they have been considered as unrelated to oxygen homeostasis, and resulting from abnormal cell function. Our results suggest that they respond to the same underlying mechanisms that maintain oxygen homeostasis in normal cells. The fact that WE involves transcriptional regulation and may require several hours to appear [71] is coherent with a HIFs-mediated response to an increase in the metabolic activity of tumor cells.

Although many attempts have been made to account for the WE in the literature [91], this is, to the best of our knowledge, the first explanation of this effect as resulting from HIFs homeostatic regulation.

The redefinition of HIFs’ homeostatic role presented in this work marks a significant departure from the prevailing paradigm, and opens new avenues for understanding the interplay between oxygen homeostasis and cellular metabolism. The proposed conceptual model sheds light on HIFs’ preventive rather than reactive function, suggesting a fundamental revision of our perception of intracellular oxygen regulation. The interest of this new approach is not merely theoretical but carries implications for therapeutic interventions targeting oxygen-responsive pathways.

## Code availability

The numerical simulations shown in Figs.3-5 have been performed using Wolfram Mathematica. The code used in these simulations is available at the Notebook Archive (https://notebookarchive.org/2024-02-cvllu1f). A PDF version of this code is included in Supplementary Material B.

## Acknowledgments

Cr.F.A. was partially supported by the MINECO grant PID2022-138187OB-I00.

