## Supplementary Material for "Redefining the role of Hypoxia-inducible factors (HIFs) in oxygen homeostasis"

### **Supplementary Information for:** Redefining the role of Hypoxia-inducible factors (HIFs) in oxygen homeostasis

### Supplementary Material A

In this section, we formalize the c [illegible] in Fig S1 below.

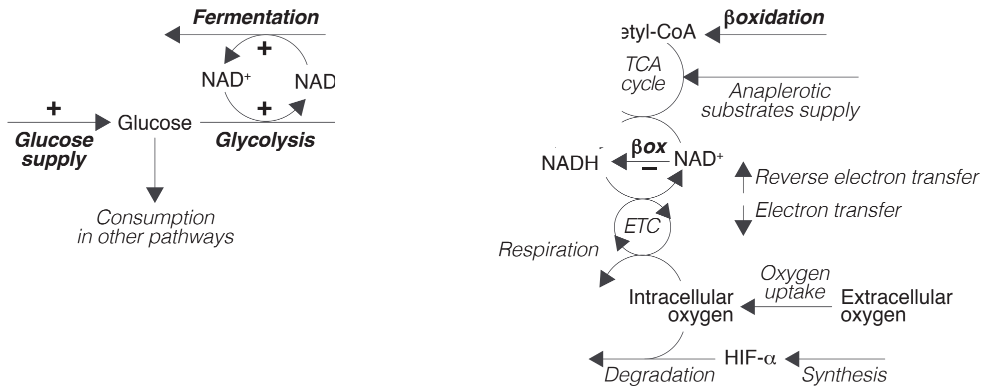


**Figure S1. Conceptual model of the physiological context of HIFs activity.** See main text for details.

This conceptual model does not aim to encompass all the details of intracellular oxygen metabolism; rather, it is conceived as a minimal system that reproduces the dynamics of oxygen uptake and consumption in relation to central catabolic processes involved in cellular energy production and regulated by HIFs. To do that, the model reduces the complexity of the reactions involved in these processes to only a few key molecules and does not take into account other elements, such as lactate or intermediate metabolites of the TCA cycle. Furthermore, it does not discriminate between cytoplasmic and mitochondrial NAD+/NADH, pyruvate, or oxygen. Instead, it condenses the metabolic network regulated by HIFs into a single compartment. The reactions included in the model can be written as:

$$\text{Glucose} + 2\ \text{NAD}^+ \xrightarrow{\gamma} 2\ \text{Pyruvate} + 2\ \text{NADH} \qquad \textit{(Glycolysis)}$$

$$\text{Pyruvate} + \text{NADH} \xrightarrow{\varphi} \text{NAD}^+ \qquad \textit{(Fermentation)}$$

$$\text{Pyruvate} + \text{NAD}^+ \xrightarrow{\delta} \text{Acetyl-CoA} + \text{NADH} \qquad \textit{(Oxidative decarboxylation)}$$

$$\text{Acetyl-CoA} + 3\ \text{NAD}^+ \xrightarrow{\tau} 3\ \text{NADH} \qquad \textit{(TCA cycle)}$$

$$\text{Fatty acids } + \text{NAD}^+ \xrightarrow{\beta} \text{Acetyl-CoA} + \text{NADH} \qquad \textit{(}\beta\textit{-Oxidation)}$$

$$\text{Anaplerotic substrates } + \text{ NAD}^+ \xrightarrow{\alpha} \text{ NADH} \qquad \textit{(TCA cycle)}$$

$$\text{ETC} + \text{NADH} \overset{\epsilon}{\rightleftarrows} \text{ETCH} + \text{NAD}^+ \qquad \textit{(Electron transfer)}$$

$$O_2 + 2\ \text{ETCH} \xrightarrow{\rho} 2\ \text{ETC}^+ \qquad \textit{(Respiration)}$$

$$O_2 + \text{HIF-}\alpha \xrightarrow{k_H} \varnothing \qquad \textit{(HIF-}\alpha\textit{ degradation)}$$

For simplicity, the model does not explicitly consider the role of $\alpha$-ketoglutarate in the PHD-mediated degradation of HIF-$\alpha$ (see Reaction 1 in the main text). All the parameters in the previous reactions take positive values. These reactions can be modeled within the framework of the law of mass action as follows:

$$\begin{cases} G'(t) = m_G + u_H H(t) - \omega G(t) - \gamma_H G(t)N(t)^2 \\ P'(t) = 2\gamma_H G(t)N(t)^2 - \delta_H P(t)N(t) - \varphi_H P(t)N_H(t) \\ A'(t) = \delta_H P(t)N(t) - \tau A(t)N(t)^3 + \beta_H m_F(t)N(t) \\ N_H'(t) = 2\gamma_H G(t)N(t)^2 + \delta_H P(t)N(t) + 3\tau A(t)N(t)^3 + \\ \qquad\qquad + \big(\alpha m_A + \beta_H m_F\big)N(t) - \varphi_H P(t)N_H(t) - \epsilon E(t)N_H(t) + \epsilon E_H(t)N(t) \\ E_H'(t) = \epsilon E(t)N_H(t) - \epsilon E_H(t)N(t) - 2\rho O_2(t)E_H(t)^2 \\ H'(t) = s_H - k_H O_2(t)H(t) \\ O_2'(t) = u_O(t) - \rho O_2(t)E_H(t)^2 - k_H O_2(t)H(t), \end{cases} \tag{1}$$

where $G(t)$, $P(t)$, $A(t)$ $O_2(t)$, and $H(t)$ are the values of glucose, pyruvate, acetyl-CoA, intracellular oxygen, and HIF-$\alpha$ at time $t$ respectively. $N(t)$ and $N_H(t)$ represent the concentration of NAD+ and NADH at time $t$ respectively, and $E(t)$ and $E_H(t)$ that of the oxidized and reduced components of the electron transport chain respectively. We have that

$$\begin{cases} N + N_H = C_{nad} \\ E + E_H = C_{etc} \end{cases} \tag{2}$$

Parameters $C_{nad}$ and $C_{etc}$ represent the capacity of the NAD+/NADH cycle and the ETC to accumulate electrons. The model inputs are denoted as $m_G$ (glucose supply), $m_A$ (anaplerotic substrates for the TCA cycle), and $m_F$ (fatty acids for $\beta$-oxidation). The rate of HIF-$\alpha$ synthesis is dentoed by $s_H$. Oxygen uptake $u_O(t)$ is given by

$$u_O(t) = \upsilon\big(E_O(t) - O_2(t)\big),$$

where $E_O(t)$ is the extracellular oxygen tension at time $t$ and $\upsilon$ a positive parameter.

HIFs-mediated regulation is modeled as:

$$\begin{cases} u_H H(t) & \textit{(Upregulation of glucose uptake)} \\ \gamma_H = \gamma + \eta H(t) & \textit{(Upregulation of glycolysis)} \\ \varphi_H = \varphi + \phi H(t) & \textit{(Upregulation of fermentation)} \\ \delta_H = \delta/\big(1 + \lambda H(t)\big) & \textit{(Dowregulation of oxidative decarboxylation)} \\ \beta_H = \beta/\big(1 + \mu H(t)\big) & \textit{(Dowregulation of } \beta\textit{-oxidation)} \end{cases} \tag{3}$$

All the model parameters used in the simulations have been chosen arbitrarily to illustrate the behavior of the model (see Supplementary Material B).

#### Supplementary Material B

This section contains the Wolfram Mathematica code used to generate Figs. 3-5 in the main text. The code is available at the Notebook Archive.

#### Model definition

In this section, we formulate a mathematical version of the following conceptual model (shown in Figure 2 in the main text):

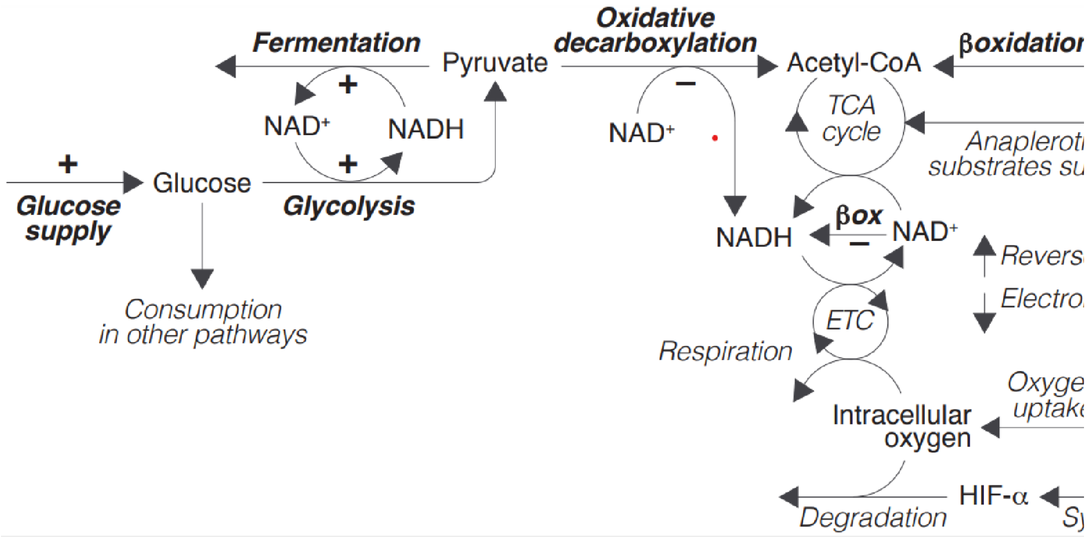


The variables and parameters of this model will be denoted as indicated below:

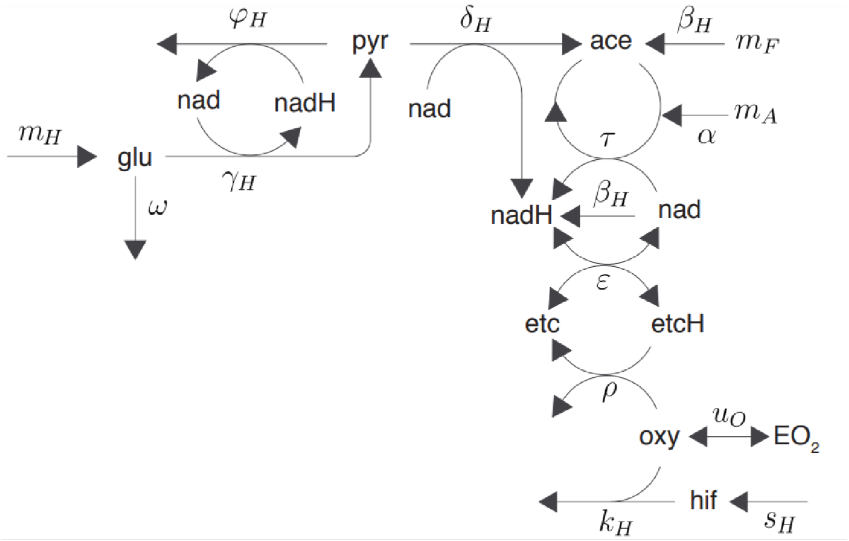


The concentration of glucose, pyruvate, acetyl-CoA, intracellular oxygen, and HIF-$\alpha$ are labeled as *glu* , *pyr*, *ace*, *oxy*, and *hif* respectively; *nad* and *nadH* represent the concentration of NAD+

and NADH respectively; and *etc* and *etcH* that of the oxidized and reduced components of the electron transport chain respectively. HIFs regulation is modeled as:

$u_H$ *(Upregulation of glucose uptake)*

$\gamma_H = \gamma + \eta H(t)$ *(Upregulation of glycolysis)*

$\varphi_H = \varphi + \phi H(t)$ *(Upregulation of fermentation)*

$\delta_H = \delta/(1 + \lambda H(t))$ *(Dowregulation of oxidative decarboxylation)*

$\beta_H = \beta/(1 + \mu H(t))$ *(Dowregulation of $\beta$-oxidation)*

The previous conceptual model can be formalized within the framework of the law of mass action as shown in Supplementary Material A. The corresponding equations can be expressed as follows:

```
Clear[model]; model[sH_, EO2_, mG_, mA_, mF_] := {
  glu'[t] == mG + uH hif[t] - ω glu[t] - (γ + η hif[t]) glu[t] × nad[t]^2,
  pyr'[t] == 2 (γ + η hif[t]) glu[t] × nad[t]^2 -
    δ pyr[t] × nad[t] / (1 + λ hif[t]) - (φ + ϕ hif[t]) pyr[t] × nadH[t],
  ace'[t] ==
   mF β nad[t] / (1 + μ hif[t]) + δ pyr[t] × nad[t] / (1 + λ hif[t]) - τ ace[t] × nad[t]^3,
  nad'[t] ==
   -3 τ ace[t] × nad[t]^3 - δ pyr[t] × nad[t] / (1 + λ hif[t]) + ϵ nadH[t] × etc[t] -
    mF β nad[t] / (1 + μ hif[t]) - mA α nad[t] + (φ + ϕ hif[t]) pyr[t] × nadH[t] -
    2 (γ + η hif[t]) glu[t] × nad[t]^2 - ϵ etcH[t] × nad[t],
  nadH'[t] == 3 τ ace[t] × nad[t]^3 + mF β nad[t] / (1 + μ hif[t]) + mA α nad[t] +
    δ pyr[t] × nad[t] / (1 + λ hif[t]) - ϵ nadH[t] × etc[t] - (φ + ϕ hif[t])
     pyr[t] × nadH[t] + 2 (γ + η hif[t]) glu[t] × nad[t]^2 + ϵ etcH[t] × nad[t],
  etc'[t] == -ϵ nadH[t] × etc[t] + 2 ρ etcH[t]^2 × oxy[t] + ϵ etcH[t] × nad[t],
  etcH'[t] == ϵ nadH[t] × etc[t] - 2 ρ etcH[t]^2 × oxy[t] - ϵ etcH[t] × nad[t],
  hif'[t] == sH - kH hif[t] × oxy[t],
  oxy'[t] == υ (EO2 - oxy[t]) - ρ etcH[t]^2 × oxy[t] - kH hif[t] × oxy[t],
  glu[0] == 0,
  pyr[0] == 1,
  ace[0] == 0,
  etc[0] == 0,
  etcH[0] == Cetc,
  nadH[0] == 0,
  nad[0] == Cnad,
  hif[0] == 0,
  oxy[0] == 1
 };

variables =
  {glu[t], pyr[t], hif[t], oxy[t], nadH[t], nad[t], ace[t], etcH[t], etcH[t]};
```

The previous equations simulate the dynamics of the conceptual model shown in Figure 2 taking as inputs the rate of HIF-$\alpha$ synthesis (*sH*), extracellular oxygen tensions (*EO2*), and the metabolic supply of glucose (*mG*), anaplerotic substrates for the TCA cycle (*mA*), and fatty acids for $\beta$-oxidation (*mF*).

```
Clear[solution];
solution[sH_, EO2_, mG_, mA_, mF_] :=
 NDSolve[model[sH, EO2, mG, mA, mF] /. parameters, variables,
  {t, 0, 200 000}, Method → {"EquationSimplification" → "Residual"}];
```

Numerical simulations range from $t = 0$ to $t = 200000$, which allows the system to stabilize and reach its steady state.

```
parameters = {uH → .5, ω → 3, γ → .8, η → .1, δ → 1.5, φ → 0.01, ϕ → 2., λ → .2, μ → .2,
   τ → .5, ϵ → 5, kH → .5, υ → 0.8, ρ → 1, α → 1, β → 1, Cnad → 100, Cetc → 500};
```

The parameters represent the rates of the metabolic pathways considered in the model, as shown in Model 1, except for *Cnad* and *Cetc*, which denote the capacity of the NAD+/NADH cycle and the electron transport chain respectively. The values of the parameters have been arbitrarily chosen to illustrate the behavior of the conceptual model outlined above.

#### Figures

##### Figure 3

In this section, we perform numerical simulations of the model for a range of values of extracellular oxygen levels (20<= *EO2*<=10000) and glucose metabolic supply (20<=*mG*<=1000), assuming that there is no HIFs-mediated regulation (i.e. that *sH*=0). In these simulations, the supply of anaplerotic substrates and fatty acids is constant: $mA = mF = 10$. These values, as well as the range of extracellular tensions and glucose supply, are chosen arbitrarily to illustrate the dynamics of the model.

```
Clear[resultsFig3];
resultsFig3 = Flatten[Table[{{EO2, mG}, (solution[0, EO2, mG, 10, 10] /.
       t → 200 000)⟦1, All, 2⟧}, {EO2, 20, 10 000, 20}, {mG, 20, 1000, 20}], 1];
```

```
Clear[data];
Table[data[variables⟦j⟧] = Table[Flatten[{resultsFig3⟦i, 1⟧,
     resultsFig3⟦i, 2, j⟧}], {i, 1, Length[resultsFig3]}], {j, 1, 9}];
```

The data are now interpolated:

```
Clear[interpolation];
Map[(interpolation[#] = Interpolation[data[#], InterpolationOrder → 2]) &,
 variables];
```

and plotted:

```
Fig3A = Plot3D[interpolation[etcH[t]][EO2, mG],
  {EO2, 20, 10 000}, {mG, 20, 1000}, MeshFunctions → {#2 &}, Mesh → 7,
  PlotPoints → 100, Ticks → None, ImageSize → 150, PlotLabel → "ECTH (%)"]
```

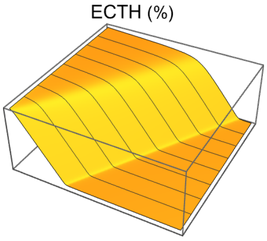

```
Fig3B = Plot3D[interpolation[nadH[t]][sO, sG],
  {sO, 20, 10000}, {sG, 20, 1000}, MeshFunctions → {#2 &}, Mesh → 7,
  PlotPoints → 100, Ticks → None, ImageSize → 150, PlotLabel → "NADH (%)"]
```

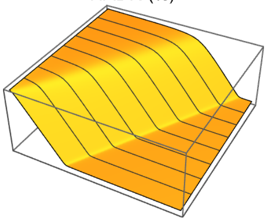


```
Fig3C = Plot3D[Log[interpolation[ace[t]][sO, sG]], {sO, 20, 9000},
  {sG, 20, 1000}, MeshFunctions → {#2 &}, Mesh → 7, Ticks → None,
  ImageSize → 150, PlotRange → All, PlotLabel → "Acetyl-CoA"]
```

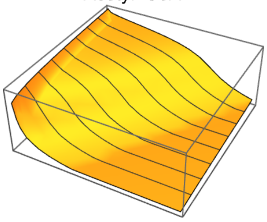


Figures 3.C and E show numerical simulations of the model for a range of values of extracellular oxygen (*EO2*) and metabolic supply (*m*) and three different rates of HIF-$\alpha$ synthesis. The metabolic supply of glucose, fatty acids, and anaplerotic substrates is assumed to increase linearly with the value of metabolic supply *m* ($mG = m$, $mA = m/20$, and $mF = m/15$). This is an arbitrary choice to illustrate the dynamics of the model. The figures show the steady-state value of NADH (%), which is calculated as 100 nadH/(nad + nadH).

```
resultsFig3C = Interpolation[Flatten[
    Table[{EO2, m, (nadH[t] /. solution[0, EO2, m, m / 20, m / 15] /. t → 200000)[[1]]},
     {EO2, 50, 9000, 50}, {m, 50, 1100, 50}], 1], InterpolationOrder → 2];
```

```
ContourPlot[resultsFig3C[EO2, m], {EO2, 50, 9000},
 {m, 50, 1100}, FrameTicks → None, ImageSize → 130, Contours → 8,
 ColorFunction → GrayLevel, AspectRatio → 1, PlotRange → All]
```

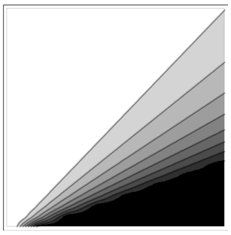

```
resultsFig3ELeft = Interpolation[Flatten[Table[
     {EO2, m, (nadH[t] /. solution[0.1, EO2, m, m / 20, m / 15] /. t → 200000)[[1]]},
     {EO2, 50, 9000, 50}, {m, 50, 1100, 50}], 1], InterpolationOrder → 2];
```

```
ContourPlot[resultsFig3ELeft[EO2, m], {EO2, 50, 9000},
 {m, 50, 1100}, FrameTicks → None, ImageSize → 130, Contours → 8,
 ColorFunction → GrayLevel, AspectRatio → 1, PlotRange → All]
```

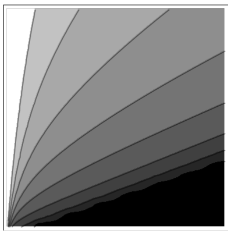

```
resultsFig3ERight = Interpolation[Flatten[Table[
     {EO2, m, (nadH[t] /. solution[2., EO2, m, m / 20, m / 15] /. t → 200 000)[[1]]},
     {EO2, 50, 9000, 50}, {m, 50, 1100, 50}], 1], InterpolationOrder → 2];
```

```
ContourPlot[resultsFig3ERight[EO2, m], {EO2, 50, 9000},
 {m, 50, 1100}, FrameTicks → None, ImageSize → 130, Contours → 8,
 ColorFunction → GrayLevel, AspectRatio → 1, PlotRange → All]
```

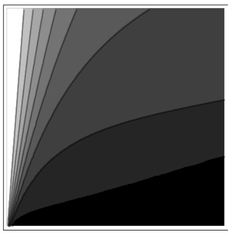

#### Figure 4

Figure 4 shows the behavior of the model for a range of values of extracellular oxygen values (50<=*EO2*<=10000).

```
extracellularOxygen = Table[EO2, {EO2, 50, 10 000, 50}];
```

Metabolic supply of glucose, anaplerotic substrates, and fatty acids is constant (*mG* = 1000, *mA* = *mF* = 10).
Numerical simulations of the model without HIFs regulation (*sH* = 0):

```
withoutHIFs = Map[solution[0, #, 1000, 10, 10] &, extracellularOxygen];
```

Numerical simulations of the model with HIFs regulation (*sH* = 10):

```
withHIFs = Map[solution[10, #, 1000, 10, 10] &, extracellularOxygen];
```

The following figures represent the dependence of the system’s steady-state on extracellular oxygen levels. Results with and without HIFs regulation are shown in orange and blue respectively.

```
Fig4A = Show[
  ListLinePlot[Transpose[
     {extracellularOxygen, Flatten[Map[hif[t] /. # /. t → 200 000 &, withHIFs]]}],
   PlotRange → {0, 50}, PlotStyle → Orange],
  ImageSize → 150,
  Ticks → {Table[EO2, {EO2, 0, 10 000, 3000}], Table[hif, {hif, 0, 50, 20}]},
  PlotLabel → "HIFα"]
```

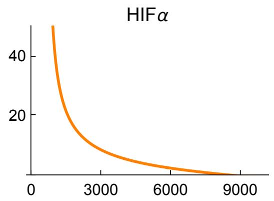


```
Fig4B = Show[
  ListLinePlot[Transpose[{extracellularOxygen,
      Flatten[Map[(φ + ϕ hif[t]) pyr[t] × nadH[t] /. parameters /. # /. t → 200 000 &,
        withHIFs]]}], PlotStyle → Orange, PlotRange → {0, 3200}],
  ListLinePlot[Transpose[{extracellularOxygen,
      Flatten[Map[(φ + ϕ hif[t]) pyr[t] × nadH[t] /. parameters /. # /. t → 200 000 &,
        withoutHIFs]]}]],
  ImageSize → 150,
  Ticks → {Table[EO2, {EO2, 0, 10 000, 3000}], Table[ferm, {ferm, 0, 3000, 1000}]},
  PlotLabel → "Fermentation"]
```

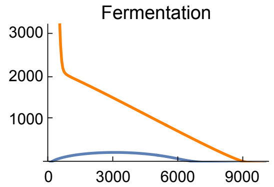


```
Fig4C = Show[ListLinePlot[Transpose[{extracellularOxygen,
      Flatten[Map[δ pyr[t] × nad[t] / (1 + λ hif[t]) /. parameters /. # /. t → 200 000 &,
        withHIFs]]}], PlotStyle → Orange],
  ListLinePlot[Transpose[{extracellularOxygen,
      Flatten[Map[δ pyr[t] × nad[t] / (1 + λ hif[t]) /. parameters /. # /. t → 200 000 &,
        withoutHIFs]]}]],
  ImageSize → 150,
  Ticks → {Table[EO2, {EO2, 0, 10 000, 3000}], Table[oxDec, {oxDec, 0, 2000, 400}]},
  PlotLabel → "Oxidative\ndecarboxylation"]
```

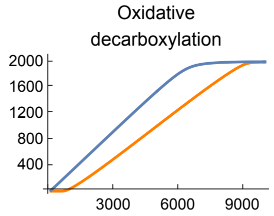

```
Fig4D = Show[
  ListLinePlot[Transpose[{extracellularOxygen, Flatten[
      Map[γ glu[t] × nad[t]^2 /. parameters /. # /. t → 200 000 &, withoutHIFs]]}]],
  ListLinePlot[Transpose[{extracellularOxygen,
     Flatten[Map[(γ + η hif[t]) glu[t] × nad[t]^2 /. parameters /. # /. t → 200 000 &,
       withHIFs]]}],
   PlotStyle → Orange, PlotRange → {0, 1500}],
  ImageSize → 150, PlotRange → {0, 1500},
  Ticks → {Table[EO2, {EO2, 0, 10 000, 3000}], Table[gly, {gly, 0, 1500, 500}]},
  PlotLabel → "Glycolysis"]
```

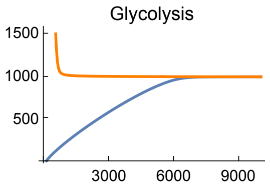


```
Fig4E = Show[
  ListLinePlot[Transpose[{extracellularOxygen,
     Flatten[Map[ρ etcH[t]^2 × oxy[t] /. parameters /. # /. t → 200 000 &,
       withHIFs]]}], PlotStyle → Orange],
  ListLinePlot[Transpose[{extracellularOxygen, Flatten[Map[
      ρ etcH[t]^2 × oxy[t] /. parameters /. # /. t → 200 000 &, withoutHIFs]]}]],
  ImageSize → 150, Ticks →
   {Table[EO2, {EO2, 0, 10 000, 3000}], Table[resp, {resp, 1000, 7000, 2000}]},
  PlotRange → {0, 7000}, PlotLabel → "Respiration"]
```

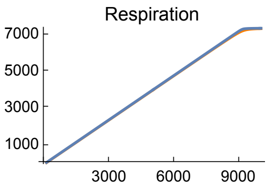


```
Fig4F = Show[
  ListLinePlot[Transpose[{extracellularOxygen,
     Flatten[Map[etcH[t] / 5 /. # /. t → 200 000 &, withHIFs]]}],
   PlotStyle → Orange, PlotRange → All],
  ListLinePlot[Transpose[{extracellularOxygen, Flatten[
      Map[etcH[t] / 5 /. # /. t → 200 000 &, withoutHIFs]]}], PlotRange → All],
  ImageSize → 150, PlotRange → {0, 100}, AxesOrigin → {0, 0},
  Ticks → {Table[EO2, {EO2, 0, 10 000, 3000}], Table[etcH, {etcH, 0, 100, 20}]},
  PlotLabel → "ETCH(%)"]
```

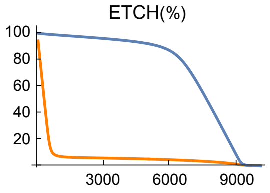

```
Fig4G = Show[
  ListLinePlot[Transpose[
     {extracellularOxygen, Flatten[Map[nadH[t] /. # /. t → 200 000 &, withHIFs]]}],
   PlotStyle → Orange, PlotRange → All],
  ListLinePlot[Transpose[{extracellularOxygen,
      Flatten[Map[nadH[t] /. # /. t → 200 000 &, withoutHIFs]]}]],
  ImageSize → 150, PlotRange → {0, 100},
  Ticks → {Table[EO2, {EO2, 0, 10 000, 3000}], Table[NADH, {NADH, 0, 200, 20}]},
  PlotLabel → "NADH(%)"]
```

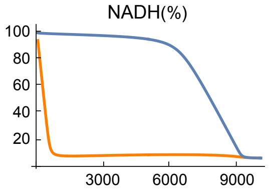


```
Fig4H = Show[
  ListLinePlot[Transpose[{extracellularOxygen,
      Flatten[Map[ace[t] /. # /. t → 200 000 &, withHIFs]]}], PlotStyle → Orange],
  ListLinePlot[Transpose[{extracellularOxygen,
      Flatten[Map[ace[t] /. # /. t → 200 000 &, withoutHIFs]]}], PlotRange → All],
  ImageSize → 150, PlotRange → {0, .03},
  Ticks → {Table[EO2, {EO2, 0, 10 000, 3000}], Table[ace, {ace, 0, .03, .01}]},
  PlotLabel → "Acetyl-CoA"]
```

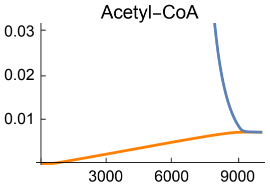


Fig4I represents the dependence of respiration on the entry of electrons in the aerobic pathway. Results with and without HIFs regulation are shown in orange and blue respectively.

```
Fig4I = Show[ListLinePlot[
   Transpose[{Flatten[Map[(δ pyr[t] × nad[t] / (1 + λ hif[t]) + τ ace[t] × nad[t] ^ 3) /.
           parameters /. # /. t → 200 000 &, withHIFs]], Flatten[
      Map[(ρ etcH[t] ^ 2 × oxy[t]) /. parameters /. # /. t → 200 000 &, withHIFs]]}],
   PlotStyle → Orange, PlotRange → All],
  ListLinePlot[
   Transpose[{Flatten[Map[(δ pyr[t] × nad[t] / (1 + λ hif[t]) + τ ace[t] × nad[t] ^ 3) /.
           parameters /. # /. t → 200 000 &, withoutHIFs]], Flatten[Map[
       (ρ etcH[t] ^ 2 × oxy[t]) /. parameters /. # /. t → 200 000 &, withoutHIFs]]}]],
  ImageSize → 250, PlotRange → All,
  Ticks → {Table[aerobic, {aerobic, 0, 40 000, 1500}],
    Table[resp, {resp, 0, 6000, 2000}]},
  AxesLabel → {"OxDec+TCA cycle", "Respiration"}]
```

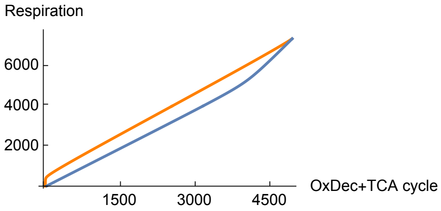


The following figures represent the dependence of fermentation and glycolysis on the rate of HIF-$\alpha$ synthesis:

```
Clear[resultsFigs4JK];
resultsFigs4JK = Table[solution[sH, 400, 1000, 10, 10], {sH, 0, 20, .05}];
```

```
Fig4J = Show[ListLinePlot[Transpose[{Table[sH, {sH, 0, 20, .05}],
     Flatten[Map[(φ + ϕ hif[t]) pyr[t] × nadH[t] /. parameters /. # /. t → 200 000 &,
       resultsFigs4JK]]}], PlotRange → All, PlotStyle → Orange],
  ImageSize → 150, PlotRange → {0, 8000},
  Ticks → {Table[sH, {sH, 0, 20, 5}], Table[ferm, {ferm, 0, 25 000, 2000}]},
  PlotLabel → "Fermentation"]
```

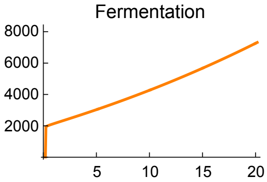

```
Fig4K = Show[ListLinePlot[Transpose[{Table[sH, {sH, 0, 20, .05}],
     Flatten[Map[(γ + η hif[t]) glu[t] × nad[t]^2 /. parameters /. # /. t → 200 000 &,
       resultsFigs4JK]]}], PlotRange → All, PlotStyle → Orange],
  ImageSize → 150, PlotRange → {0, 4000},
  Ticks → {Table[sH, {sH, 0, 20, 5}], Table[gly, {gly, 0, 4000, 1000}]},
  PlotLabel → "Glycolysis"]
```

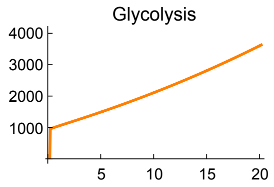


#### Figure 5

Figure 5 shows the response of the model to changes in metabolic supply (*m*). The input of glucose, fatty acids, and anaplerotic substrates into the system increases linearly with the value of metabolic supply *m* ($mG = m$, $mA = m/20$, and $mF = m/15$). Extracellular oxygen tension is constant ($EO2$=4000).

```
metabolicSupply = Table[m, {m, 0, 3000, 2}];
```

Numerical simulations of the model without HIFs regulation ($sH = 0$):

```
withoutHIFs = Map[solution[0, 4000, #, # / 20, # / 15] &, metabolicSupply];
```

Numerical simulations of the model with HIFs regulation ($sH = 10$):

```
withHIFs = Map[solution[10, 4000, #, # / 20, # / 15] &, metabolicSupply];
```

The following figures show the dependence of the system's steady-state on metabolic supply. Results with and without HIFs regulation are shown in orange and blue respectively.

```
Fig5A = Show[
  ListLinePlot[Transpose[{metabolicSupply,
     Flatten[Map[hif[t] /. parameters /. # /. t → 200 000 &, withHIFs]]}],
   PlotRange → All, PlotStyle → Orange], ImageSize → 150, PlotRange → {0, 500},
  Ticks → {Table[m, {m, 0, 3000, 1000}], Table[hif, {hif, 0, 500, 150}]},
  PlotLabel → "HIF-α"]
```

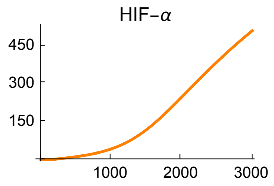

```
Fig5B = Show[
  ListLinePlot[Transpose[{metabolicSupply,
      Flatten[Map[etcH[t] / 5 /. parameters /. # /. t → 200 000 &, withHIFs]]}],
   PlotRange → All, PlotStyle → Orange],
  ListLinePlot[Transpose[{metabolicSupply,
      Flatten[Map[etcH[t] / 5 /. parameters /. # /. t → 200 000 &, withoutHIFs]]}],
   PlotRange → All], ImageSize → 150, PlotRange → {0, 100},
  Ticks → {Table[m, {m, 0, 3000, 1000}], Table[NADH, {NADH, 0, 100, 20}]},
  PlotLabel → "NADH (%)"]
```

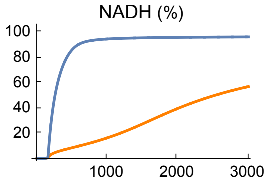


```
Fig5C = Show[ListLinePlot[Transpose[{metabolicSupply,
      Flatten[Map[δ pyr[t] × nad[t] / (1 + λ hif[t]) /. parameters /. # /. t → 200 000 &,
        withHIFs]]}], PlotStyle → Orange,
   PlotRange → All], ListLinePlot[Transpose[{metabolicSupply,
      Flatten[Map[δ pyr[t] × nad[t] / (1 + λ hif[t]) /. parameters /. # /. t → 200 000 &,
        withoutHIFs]]}], PlotRange → All], ImageSize → 150, PlotRange → {0, 1100},
  Ticks → {Table[m, {m, 0, 3000, 1000}], Table[oxDec, {oxDec, 0, 4000, 250}]},
  PlotLabel → "Oxidative\ndecarboxylation"]
```

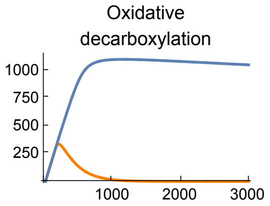


```
Fig5D = Show[
  ListLinePlot[Transpose[{metabolicSupply, Flatten[
       Map[τ ace[t] × nad[t] ^3 /. parameters /. # /. t → 200 000 &, withHIFs]]}],
   PlotStyle → Orange, PlotRange → All], ListLinePlot[
   Transpose[{metabolicSupply, Flatten[
       Map[τ ace[t] × nad[t] ^3 /. parameters /. # /. t → 200 000 &, withoutHIFs]]}],
   PlotRange → All], ImageSize → 150, Ticks → {Table[m, {m, 0, 3000, 1000}],
    Table[tca, {tca, 0, 2000, 400}]}, PlotLabel → "TCA cycle"]
```

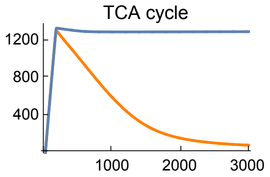

```
Fig5E = Show[Show[ListLinePlot[Transpose[{metabolicSupply, Flatten[
        Map[γ glu[t] × nad[t]^2 /. parameters /. # /. t → 200 000 &, withHIFs]]}]],
    ImageSize → 150, PlotRange → All], ListLinePlot[Transpose[{metabolicSupply,
      Flatten[Map[(γ + η hif[t]) glu[t] × nad[t]^2 /. parameters /. # /. t → 200 000 &,
        withoutHIFs]]}], PlotStyle → Orange], ImageSize → 150,
  PlotRange → {0, 800}, Ticks → {Table[m, {m, 0, 3000, 1000}],
    Table[gly, {gly, 0, 800, 200}]}, PlotLabel → "Glycolysis"]
```

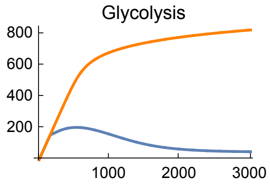


```
Fig5F = Show[ListLinePlot[Transpose[{metabolicSupply,
      Flatten[Map[ϕ hif[t] × pyr[t] × nadH[t] + φ pyr[t] × nadH[t] /. parameters /. # /.
          t → 200 000 &, withHIFs]]}], PlotRange → All, PlotStyle → Orange],
  ListLinePlot[Transpose[{metabolicSupply,
      Flatten[Map[ϕ hif[t] × pyr[t] × nadH[t] + φ pyr[t] × nadH[t] /. parameters /. # /.
          t → 200 000 &, withoutHIFs]]}],
   PlotRange → All], ImageSize → 150, PlotRange → {0, 7000},
  Ticks → {Table[m, {m, 0, 3000, 1000}], Table[fer, {fer, 0, 7000, 2000}]},
  PlotLabel → "Fermentation"]
```

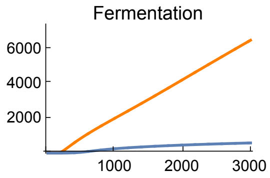


Figure 5.G shows lactate production as a function of metabolic supply for three different values of extracellular oxygen tension (*EO2* = 1000, 4000, and 20000). These values are chosen arbitrarily to represent low, intermediate, and high values of oxygen availability.

```
withHIFsHypoxia = Map[solution[10, 1000, #, # / 20, # / 15] &, metabolicSupply];
```

```
withHIFsNormoxia = Map[solution[10, 20 000, #, # / 20, # / 15] &, metabolicSupply];
```

```
Fig5G = Show[ListLinePlot[Transpose[{metabolicSupply,
      Flatten[Map[ϕ hif[t] × pyr[t] × nadH[t] + φ pyr[t] × nadH[t] /. parameters /. # /.
          t → 200 000 &, withHIFsHypoxia]]}], PlotRange → All, PlotStyle → Green],
   ListLinePlot[Transpose[{metabolicSupply,
      Flatten[Map[ϕ hif[t] × pyr[t] × nadH[t] + φ pyr[t] × nadH[t] /. parameters /. # /.
          t → 200 000 &, withHIFs]]}], PlotRange → All, PlotStyle → Orange],
   ListLinePlot[Transpose[{metabolicSupply,
      Flatten[Map[ϕ hif[t] × pyr[t] × nadH[t] + φ pyr[t] × nadH[t] /. parameters /. # /.
          t → 200 000 &, withHIFsNormoxia]]}],
    PlotRange → All], ImageSize → 150, PlotRange → {0, 11 000},
   Ticks → {Table[m, {m, 0, 3000, 1000}], Table[fer, {fer, 0, 12 000, 3000}]},
   PlotLabel → "Lactate production"]
```

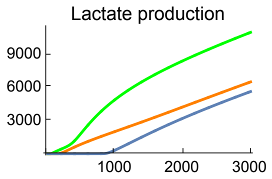


Figure 5.H shows the changes in lactate production with the rate of HIF-$\alpha$ synthesis ($sH$) for three values of extracellular oxygen tensions ($EO2$ = 1000, 4000, and 20000). These values are chosen arbitrarily to represent low, intermediate, and high values of oxygen availability.

```
hypoxia = Table[solution[sH, 1000, 2000, 2000 / 20, 2000 / 15], {sH, 0, 20, .05}];

physioxia = Table[solution[sH, 4000, 2000, 2000 / 20, 2000 / 15], {sH, 0, 20, .05}];

normoxia = Table[solution[sH, 20 000, 2000, 2000 / 20, 2000 / 15], {sH, 0, 20, .05}];

Fig5H = Show[ListLinePlot[Transpose[{Table[sH, {sH, 0, 20, .05}],
      Flatten[Map[(ϕ hif[t] + φ) pyr[t] × nadH[t] /. parameters /. # /. t → 200 000 &,
        hypoxia]]}], PlotStyle → Orange, PlotRange → All],
   ListLinePlot[Transpose[{Table[sH, {sH, 0, 20, .05}],
      Flatten[Map[(ϕ hif[t] + φ) pyr[t] × nadH[t] /. parameters /. # /. t → 200 000 &,
        physioxia]]}], PlotRange → All],
   ListLinePlot[Transpose[{Table[sH, {sH, 0, 20, .05}],
      Flatten[Map[(ϕ hif[t] + φ) pyr[t] × nadH[t] /. parameters /. # /. t → 200 000 &,
        normoxia]]}], PlotStyle → Red, PlotRange → All], ImageSize → 150,
   Ticks → {Table[sH, {sH, 0, 20, 4}], Table[sO, {sO, 0, 12 000, 4000}]},
   PlotRange → All, PlotLabel → "Lactate production"]
```

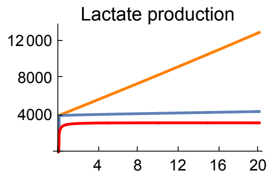
